# Differential Impact of Lymphocytes on Radiation Response in Autochthonous versus Transplant Sarcomas in Syngeneic Mice

**DOI:** 10.1101/2025.10.05.680584

**Authors:** Yvonne M. Mowery, Aastha Sobti, Alex M. Bassil, Amy J. Wisdom, Collin L. Kent, Chang Su, Jonathan Himes, Eric S. Xu, Nerissa T. Williams, Lixia Luo, David G. Kirsch

## Abstract

Preclinical studies in transplant tumor models showing high cure rates with combined radiotherapy (RT) and immunotherapy have rarely translated to clinical success, and studies in autochthonous tumor models are limited. We hypothesized that lymphocytes differentially affect RT response in transplant versus autochthonous tumor models. Here, we compared tumor onset and growth delay after 0 or 20 Gy for autochthonous versus transplant soft tissue sarcomas in *Rag2*^−/−^ mice lacking an adaptive immune system versus immune-intact *Rag2*^+/−^ mice. While time to tumor onset did not differ between *Rag2*^−/−^ and *Rag2*^+/−^ mice in the autochthonous model, transplant tumor onset was significantly slower in immunocompetent mice. No transplant tumors in *Rag2*^−/−^ mice or autochthonous tumors were cured by RT, whereas 30.4% of *Rag2*^+/−^ mice with transplant tumors were cured. These results highlight the importance of including autochthonous tumors as complementary model systems to study the interplay between the immune system and RT response.

---

Potential synergy between radiotherapy (RT) and immunotherapy has garnered attention due to studies in transplant tumor models showing high cure rates and abscopal responses with combinations of RT and immune checkpoint inhibitors (ICI) [1–6]. However, translating promising preclinical findings into clinical success has been challenging, and numerous clinical trials testing ICI with RT failed to demonstrate significant benefit [7–10]. This discrepancy between preclinical and clinical findings underscores the importance of reassessing factors that shape immune-RT interactions in various tumor models.

Helen Stone’s seminal work in 1979 demonstrated that immune suppression promoted RT resistance in a transplant fibrosarcoma model, whereas immune stimulation sensitized tumors to RT [1]. Notably, injecting cancer cells into a syngeneic host can activate the immune system, leading to immune recognition and response against the transplanted tumor [11–13]. Autochthonous tumors, arising over weeks to months under immune surveillance, may exhibit distinct immune dynamics that influence therapeutic response [14]. Here, we evaluate the role of lymphocytes in immune surveillance and RT response in transplant versus autochthonous tumors from a high mutational load soft tissue sarcoma mouse model (~2000 nonsynonymous somatic mutations/ tumor) [15]. We hypothesized that lymphocytes have a greater effect on immune surveillance and RT response in transplant tumors than autochthonous tumors.

Mouse studies were approved by the Institutional Animal Care and Use Committee. Primary and transplant p53/3-methylcholanthrene (MCA) sarcomas were generated in male and female mice aged 6-10 weeks as previously described [14–16]. Autochthonous p53/MCA sarcomas (**Figure 1A**) were induced by intramuscular (IM) injection of adenovirus-expressing Cre recombinase (Adeno-Cre, University of Iowa Viral Vector Core) and 300 μg MCA (MilliporeSigma) in sesame oil (6 μg/μl) into the gastrocnemius muscle of *Rag2*^−/−^; *Trp53*^*fl/fl*^ or *Rag2*^+/−^; *Trp53*^*fl/fl*^ mice (immune-intact littermate controls). Transplant sarcomas (**Figure 1B**) utilized a p53/MCA cell line generated in a C57BL/6J mouse by IM injection of adenovirus expressing Cas9 endonuclease and sgRNA targeting *Trp53* (sgp53: GATGGTAAGGATAGGTCGG) (Adeno-sgp53-Cas9; Viraquest) [14, 16, 17]. Tumors were induced by IM injection of 50,000 cells (100□µL, 1:1 DMEM (Gibco): Matrigel (Corning)) into the gastrocnemius muscle of *Rag2*^−/−^ (Jackson Lab, strain 008309) or *Rag2*^+/−^ mice on a C57BL/6J background.

**Figure 1.**
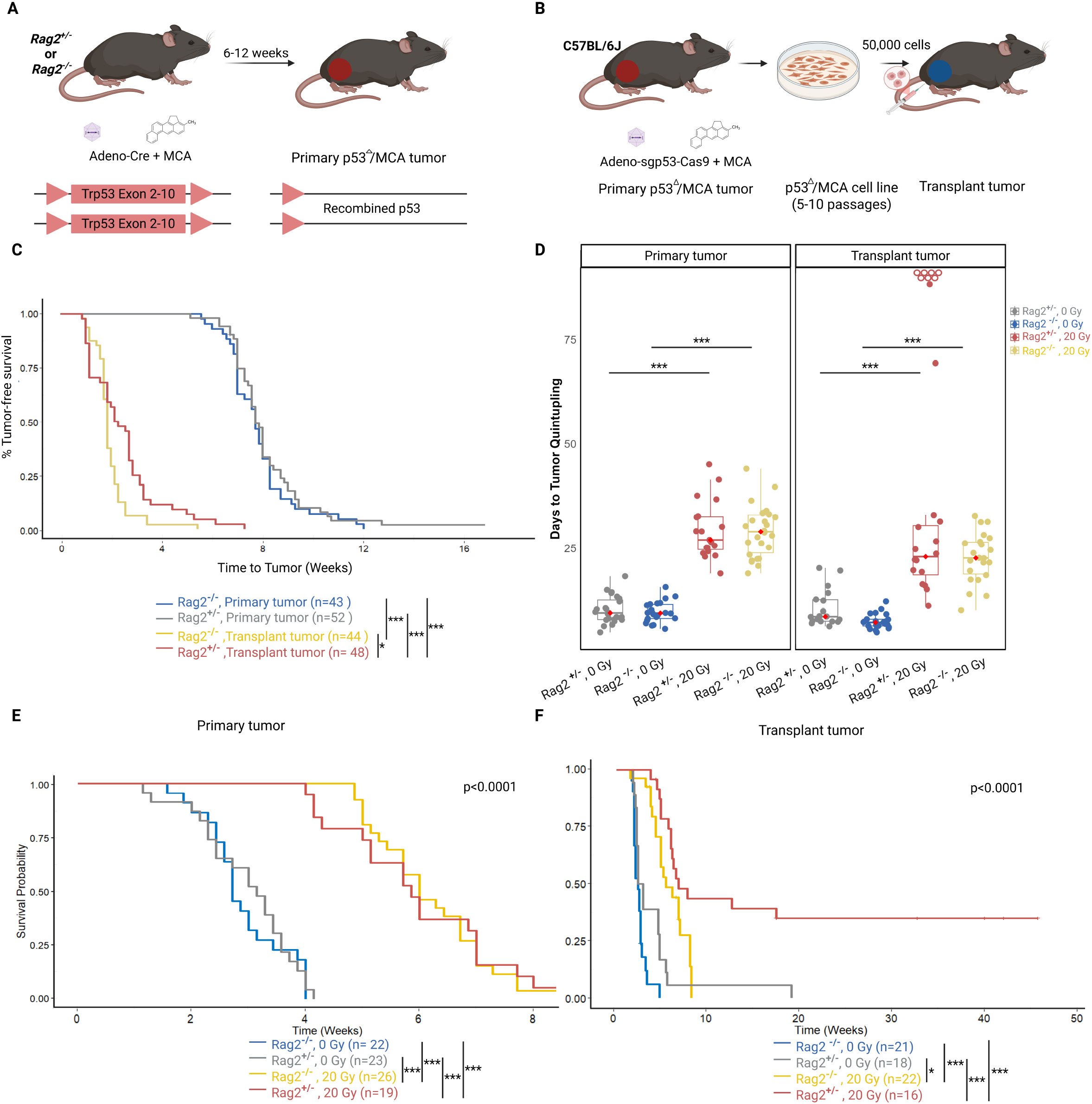
Differential Effect of Lymphocyte Deficiency on Tumorigenesis, Growth Kinetics, and Radiation Response in Autochthonous versus Transplant Sarcomas. **A**) Autochthonous p53/MCA sarcomas were induced by IM injection of Adeno-Cre and MCA in *Rag2*^−/−^; *Trp53*^*fl/fl*^ mice or *Rag2*^*+/*^; *Trp53*^*fl/fl*^ mice. **B.** Transplant p53/MCA sarcomas were induced in *Rag2*^+/−^ and *Rag2*^−/−^ mice (C57BL/6J background) by IM injection of a p53/MCA cell line generated from a tumor induced by IM injection of Adeno-sgp53-Cas9 and MCA in a C57BL/6J mouse. **C**. Kaplan-Meier plots show the time to tumor development (defined as tumor volume >70 mm^3^) for autochthonous and transplant tumors in *Rag2*^−/−^ mice or *Rag2*^*+l-*^ mice. Pairwise comparisons using log-rank test with Bonferroni correction between genotypes for each model were performed. **D**. Box plots show time to tumor volume quintupling in days for autochthonous (left) and transplant (right) tumors in *Rag2*^+/−^ and *Rag2*^−/−^ mice after treatment with 0 or 20 Gy (ANOVA and Tukey’s post-hoc tests). Seven out of 23 *Rag2*^+/−^ mice with transplant tumors were cured and are shown in the plot as unfilled circles in red. The boxes represent the interquartile range (IQR), with whiskers extending to 1.5 times the IQR from the lower (Q1) and upper (Q3) quartiles. The red dots denote median values for each variable. Jitter points demonstrate individual observations, highlighting the distribution in each group. **E**. Kaplan-Meier plots show overall survival after treatment with 0 or 20 Gy in *Rag2*^+/−^ and *Rag2*^−/−^ mice with autochthonous tumors. Pairwise comparison by the log-rank test with Bonferroni correction between genotypes for each treatment group is shown. **F**. Kaplan-Meier plots show overall survival time after treatment with 0 or 20 Gy in *Rag2*^+/−^ and *Rag2*^−/−^ mice with transplant tumors. Adjusted p-values after Bonferroni correction: *, p < 0.05; **, p < 0.01; ***, p < 0.001.

When tumors reached 70-120 mm^3^, mice were randomized to receive 0 or 20 Gy (single fraction) (Precision Xrad-225Cx). Parallel-opposed anterior/posterior 40×40 mm^2^ fields (225□kVp, 13□mA X-rays, prescribed to midplane at isocenter, average dose rate 300 cGy/min) [14]. Statistical analysis included time to tumor volume quintupling (TTQ), time to tumor onset, and overall survival after treatment. Statistical analyses were conducted using RStudio (Boston, MA, USA) (R Version 4.4.1, Vienna, Austria), with p < 0.05 considered significant.

In the autochthonous model, lymphocyte deficiency did not significantly affect tumorigenesis, tumor growth kinetics, or RT response. Median time to tumor development was 54 days in *Rag2*^+/−^ and *Rag2*^−/−^ mice (**Figure 1C**). For unirradiated tumors, median TTQ was 9.3 days in *Rag2*^+/−^ and *Rag2*^−/−^ mice. RT delayed tumor growth relative to unirradiated tumors, but median TTQ did not differ significantly in the presence or absence of mature lymphocytes (median 26.9 days for *Rag2*^+/−^ vs. 28.9 days for *Rag2*^−/−^ mice; *p* = 1) (**Figure 1D**). Additionally, no mice with autochthonous tumors were cured by 20 Gy. Survival was comparable between both genotypes for the unirradiated (median survival 22 days in *Rag2*^+/−^ vs. 19 days in *Rag2*^−/−^ mice; *p* = 0.1) and 20 Gy groups (median survival 41 days for *Rag2*^+/−^ vs. 42 days for *Rag2*^−/−^ mice; *p* = 1) (**Figure 1E**). These data show that lymphocytes do not significantly influence tumor development, growth, or RT response in the autochthonous p53/MCA sarcoma model.

By contrast, the transplant p53/MCA model revealed distinct differences in tumor onset and RT response based on the presence of mature lymphocytes. Median time to tumor onset was significantly longer in *Rag2*^+/−^ mice compared to immunodeficient *Rag2*^−/−^ mice (15.5 vs. 13 days, respectively; *p* = 0.006), indicating that lymphocytes affect early establishment of transplant sarcomas, but not autochthonous p53/MCA tumors (**Figure 1C**). Unirradiated tumor growth was faster in mice lacking lymphocytes, as evidenced by significantly shorter median TTQ of 7.2 days in *Rag2*^−/−^ mice versus 8.5 days in *Rag2*^+/−^ mice (*p* = 0.01) (**Figure 1D**). Survival was longer in *Rag2*^+/−^ mice compared to *Rag2*^−/−^ mice with unirradiated transplant tumors, but this was not statistically significant (median survival 19 vs. 17 days, *p* = 0.19). While no *Rag2*^−/−^ mice with transplant tumors were cured with RT, irradiated tumors were eliminated in a substantial proportion of *Rag2*^+/−^ mice (7/23, 30.4%). No tumors developed after tumor re-challenge with the same p53/MCA cell line injected into the contralateral leg of cured mice (n=7), demonstrating immunologic memory. Survival after 20 Gy to transplant tumors was significantly longer in *Rag2*^+/−^ compared to *Rag2*^−/−^ mice (median survival 49 vs. 39 days; *p* = 0.03) (**Figure 1F**). These findings underscore the critical role of lymphocytes in RT response in the transplant sarcoma model, highlighting the importance of adaptive immunity in achieving tumor cure after RT.

This study demonstrates a differential role of lymphocytes in RT response for transplant and autochthonous sarcomas from the p53/MCA model system. These findings highlight the complexity of tumor-immune interactions in preclinical studies and their implications for therapeutic responses. Our results in the transplant p53/MCA sarcoma model mirror the historic study by Stone et al. showing the immune system’s importance in RT-induced control of transplant tumors. However, these findings do not necessarily apply to autochthonous tumors that gradually develop under immune surveillance, similar to spontaneous tumors in patients. In the autochthonous p53/MCA model, the presence or absence of functional lymphocytes did not significantly impact tumor onset, growth kinetics, or RT response. This differential role of lymphocytes in response to RT across models highlights the need for tailored treatment approaches based on the specific immune context of each tumor [14].

We previously characterized the distinct immune landscapes in autochthonous and transplant p53/MCA sarcomas and demonstrated their differential responses to RT and anti-PD-1 therapy [14]. Untreated transplant tumors were enriched for CD8^+^ T cells and M2 macrophages, resembling highly inflamed human sarcomas that are more likely to respond to PD-1 checkpoint blockade [12]. Autochthonous sarcomas demonstrated immunoediting, reduced neoantigen expression, and tumor-specific immune tolerance, and their immune profile resembled poorly immunogenic human undifferentiated pleomorphic sarcomas, which are less likely to respond to PD-1 inhibition [12, 14]. While autochthonous tumors were resistant to RT and anti-PD-1 therapy, mice with transplant tumors were cured by this combination. Subsequent T cell depletion experiments demonstrated that response to RT and anti-PD-1 therapy in the transplant model was dependent on CD8^+^ T cells. The current study similarly demonstrated a role for lymphocytes in RT response in the transplant model, with a subset of *Rag2*^+/−^ mice bearing transplant p53/MCA tumors cured after 20 Gy and immune to rechallenge. By contrast, no transplant tumors in *Rag2*^−/−^ mice or autochthonous tumors in immunocompetent or immunodeficient mice were cured by RT.

The artificial nature of immune-mediated response to RT in transplant models may be attributed to several factors including the absence of co-evolution between tumor and host immune system, potential immune activation/priming during transplantation, and the lack of a fully developed immunosuppressive tumor microenvironment [13, 14, 18-20]. Studies indicate that initial CD8^+^ T cell-mediated immune responses to implanted tumors can be adoptively transferred to other hosts, effectively managing corresponding tumors from the same model. However, early anti-tumor CD8^+^ T cell responses can be followed by emergence of inhibitory CD4^+^ regulatory T cells, highlighting the dynamic characteristics of immune response to transplanted tumors [13, 18–20]. In immunocompetent preclinical models, a robust initial immune response can lead to spontaneous tumor regression, while the same tumors grow progressively in immunodeficient mice. This dichotomy underscores the critical role of host immune competence in modulating tumor progression and highlights the potential for immune-mediated tumor control in the context of transplantation models [21, 22]. These differential therapeutic responses emphasize the importance of studying RT effects in both transplant and autochthonous tumor models. Limitations to our study include the use of a single tumor model and high-dose single-fraction radiation regimen, potentially limiting generalizability. Furthermore, the focus on lymphocytes neglects potential contributions of other immune cell populations, such as tumor-associated macrophages and dendritic cells, which play pivotal roles in the tumor microenvironment and may significantly influence treatment outcomes [23–26]. To understand the complex interplay between radiotherapy and the immune system, future investigations should take a comprehensive approach—incorporating diverse tumor models, varying radiation fractionation schemes, and examining interactions among multiple immune cell subsets.

In conclusion, our findings highlight the importance of employing complementary preclinical model systems, including autochthonous tumors, to investigate complex interactions among tumors, radiotherapy, and the immune system. The discrepancy between promising preclinical findings and disappointing clinical outcomes with radiotherapy and immunotherapy may partly stem from shortcomings of conventional preclinical transplant models to accurately reflect human tumor biology. Autochthonous tumors may recapitulate human cancers that exhibit reduced responsiveness to RT and immunotherapy. Consequently, additional research utilizing these complementary models is important to advance therapies with increased translational potential. Focusing on effective strategies in autochthonous tumor models may help to bridge the gap between preclinical promise and clinical efficacy.

## Acknowledgements

This work was supported by NIH grants to YMM K08DE029887 and to DGK R35CA197616 and by an ASCO Young Investigator Award to YMM. The figure was assembled in Biorender.

